# Generative Haplotype Prediction Outperforms Statistical Methods for Small Variant Detection in NGS Data

**DOI:** 10.1101/2024.02.27.582327

**Authors:** Brendan O’Fallon, Ashini Bolia, Jacob Durtschi, Luobin Yang, Eric Fredrickson, Hunter Best

## Abstract

Detection of germline variants in next-generation sequencing data is an essential component of modern genomics analysis. Variant detection tools typically rely on statistical algorithms such as de Bruijn graphs or Hidden Markov Models, and are often coupled with heuristic techniques and thresholds to maximize accuracy. Here, we introduce a new approach that replaces these handcrafted statistical techniques with a single deep generative model. The model’s input is the set of reads aligning to a single genomic region, and the model produces two sets of output tokens, each representing the nucleotide sequence of a germline haplotype. Using a standard transformer-based encoder and double-decoder architecture, our model learns to construct germline haplotypes in a generative fashion identical to modern Large Language Models (LLMs). We train our model on 37 Whole Genome Sequences (WGS) from Genome-in-a-Bottle samples, and demonstrate that our method learns to produce accurate haplotypes with correct phase and genotype for all classes of small variants. We compare our method, called Jenever, to FreeBayes, GATK HaplotypeCaller, Clair3 and DeepVariant, and demonstrate that our method has superior overall accuracy compared to other methods. At *F*1-maximizing quality thresholds, our model delivers the highest sensitivity, precision, and the fewest genotyping errors for insertion and deletion variants. For single nucleotide variants our model demonstrates the highest sensitivity but at somewhat lower precision, and achieves the highest overall *F*1 score among all callers we tested.

## 1 Introduction

Identifying the DNA sequence variants in a sample is one of the most common and important tasks in modern genomics. Tools that aim to identify variants must overcome several challenges to produce an accurate set of variant calls. For instance, the sequenced reads themselves may contain errors in the form of miscalled bases or PCR-induced repeat length errors, and the tools used to align reads to the reference genome may produce incorrect mappings (e.g. Li et al. 2014, Goldfeder et al. 2016). In addition, large or complex sequence variants may be obscured by limitations in the read alignment algorithm, and require a sophisticated assembly of reads to identify. Reconciling artifacts produced during lab preparation procedures, sequencing, basecalling, and read alignment is a significant technical challenge, and modern variant detection tools employ a variety of statistical and heuristic techniques to achieve high precision and sensitivity (e.g. De Pristo et al. 2011, Nielson et al. 2011).

All variant discovery tools must address two challenges. First, callers must identify potential alleles in a region. Then, given a set of potential alleles, they must assess the likelihood of each candidate (or pairs of candidates in the diploid case) to determine which should be included in the caller output. Early callers, such as samtools / mpileup (Li et al. 2009) and the UnifiedGenotyper tool from the Genome Analysis ToolKit (GATK, De Pristo et al. 2011) rely on the read aligner to generate candidate alleles, and utilize several ad-hoc heuristics and thresholds to determine which alleles are most likely (e.g. Nielsen et al. 2011). Later tools (e.g. Poplin et al. 2018, Kim et al. 2018, Cooke et al. 2021) incorporated local re-assembly of reads in each region of interest, resulting in a significant improvement in the sensitivity and precision of variant calls. For instance, the HaplotypeCaller tool identifies candidate haplotypes by constructing de Bruijn graphs from k-mers present in the reads, and assesses haplotype likelihoods with a pair Hidden Markov Model (pair-HMM). The de Bruijn graph / pair-HMM approach has been adopted by multiple other top-performing tools, including DeepVariant (Poplin et al. 2018) and DRAGEN (Behera et al. 2024)

More recent tools have incorporated elements of deep learning into the allele likelihood calculation or allele generation steps. DeepVariant (Poplin et al. 2018) uses a statistical method based on the HaplotypeCaller approach to identify candidates, but adds a Convolutional Neural Net (CNN) to classify variants as true or false positive detections. HELLO (Ramachandran et al. 2020), designed to work on hybrid short- and long-read datasets, employs a mixture-of-experts approach with separate 1-dimensional convolutions across the read and position dimensions of the input. Clair (Luo et al. 2019, Luo et al. 2020), uses a novel multitask deep learning approach to predict several properties of a potential variant at a given site, including zygosity and allele length. DAVI (Gupta & Saini 2020) is a unique tool where both primary read mapping and variant detection are performed using neural networks, although the variant detection component was limited to SNPs only and DAVI is unable to reconstruct more complex forms of variation.

Most current variant detection tools that utilize deep learning techniques employ CNNs as the primary architecture (e.g. Poplin et al. 2018, Gupta & Saini 2020, Luo et al. 2020). CNNs have proven successful in computer visions tasks in part due to translational invariance, which helps the model to recognize local features or patterns regardless of their absolute position in an image. In variant detectors, CNNs are often used in an image classification context, such that the primary output is a probability distribution over classes representing, for instance, possible genotypes. However, CNN architectures are less well suited to candidate allele generation, and methods have used statistical approaches (e.g. Poplin et al. 2018) to generate candidates, or more complicated multi-task approaches with separate tasks for genotype, allele length and phasing (e.g. Luo et al. 2020).

The transformer architecture (Vaswani et al. 2017) offers a more straightforward approach to allele generation than CNNs. Because generative transformer models can create arbitrarily long or complex output sequences, they do not require a separate allele generation step and can be trained end-to-end to produce the desired output. End-to-end training of the full model is promising since the hand-crafted approaches used in prior methods can be replaced by a single, unified model. Recently Baid et al (2023) used a transformer-encoder model to improve the accuracy of PacBio consensus reads, and our group described how aligned NGS reads can be tokenized in a manner suitable for input to a transformer model (O’Fallon et al. 2022), but neither project explored transformer-based decoding. To our knowledge transformer-based models of haplotype generation have not been explored previously.

Here, we describe a new approach to small variant detection in NGS data that uses a generative, transformer-based model to predict the two germline haplotypes present in the sample. The inputs to our model are the series of bases aligning to individual reference positions, and the outputs are the predicted haplotypes. Each output token of our model is a 4-mer of nucleotide bases, and multiple output tokens are generated to represent full haplotype sequences. This approach does not require any of the bespoke statistical machinery or heuristic thresholds utilized by most modern callers, and instead uses a single deep learning model to reconstruct complete haplotypes directly from aligned NGS reads.

## 2 Methods

### 2.1 Model architecture & read encoding

Our model consists of a single transformer-based encoder module connected to two transformer decoders, one for each haplotype. The encoder and decoder components are standard transformers with GeLU activations as implemented in PyTorch 2.1 (Vaswani et al. 2017, Paszke et al. 2019). Input tokens are the collection of bases that align to a given reference position, with some modifications described below. We add an additional fully connected layer prior to the transformer encoders which embeds the encoded basecalls in *d* dimensions, where *d* = 12 for the analyses here. The embedded basecalls are ‘flattened’ along the read dimension, producing an input token with size *dr* where *r* is the number of reads (*r* = 150 for all experiments reported here). Instead of the typical 1-dimensional positional encoding, we employ a 2-dimensional encoding, allowing the model to differentiate across both the token (position) and read dimensions. The 2D encoding implementation follows Wang & Jyh-Charn (2021).

Input tensors are generated by selecting a 150bp region from a set of aligned reads in BAM / CRAM format. For each region, all reads overlapping the window were obtained from the corresponding alignment file. Regions containing more than the maximum number of reads (150 for all analyses here) were downsampled to the maximum size. Reads were sorted by the reference coordinate of the first aligned base. For each aligned base in the selected reads ten features are encoded; the first four are the one-hot encoded base call, followed by base quality, two flags indicating if base ‘consumed’ reference base (i.e. was not an insertion) or consumed a sequence read base (was not a deletion), and additional flags indicating sequence read direction, clipping status, and mapping quality. No gap tokens or other special handling was performed for insertions or deletions. In addition to sequenced reads, the first row in each encoded region was the reference sequence. For this special row we inserted a base quality of 100 for every position and did not set any of the other flags. Resulting tensors had dimension [*g, r*, 10], where *g* indexes genomic positions, *r* indexes reads, and 10 is the number of features. Positions in the input not corresponding to an aligned base were all set to zero.

Our model uses two parallel transformer decoder blocks, each of which produces output tokens representing a single haplotype. Output tokens are *k*-mers of length 4, and the decoders do not share parameters. As there are four possible bases at each position both decoders have model dimension 4^4^ = 256. Because the encoder and decoders do not necessarily have equal dimensions we convert encoder logits to dimension 256 with a single linear layer. A standard 1-dimensional positional encoding is applied to the encoder logits prior to decoding (see Figure 1).

**Figure 1:**
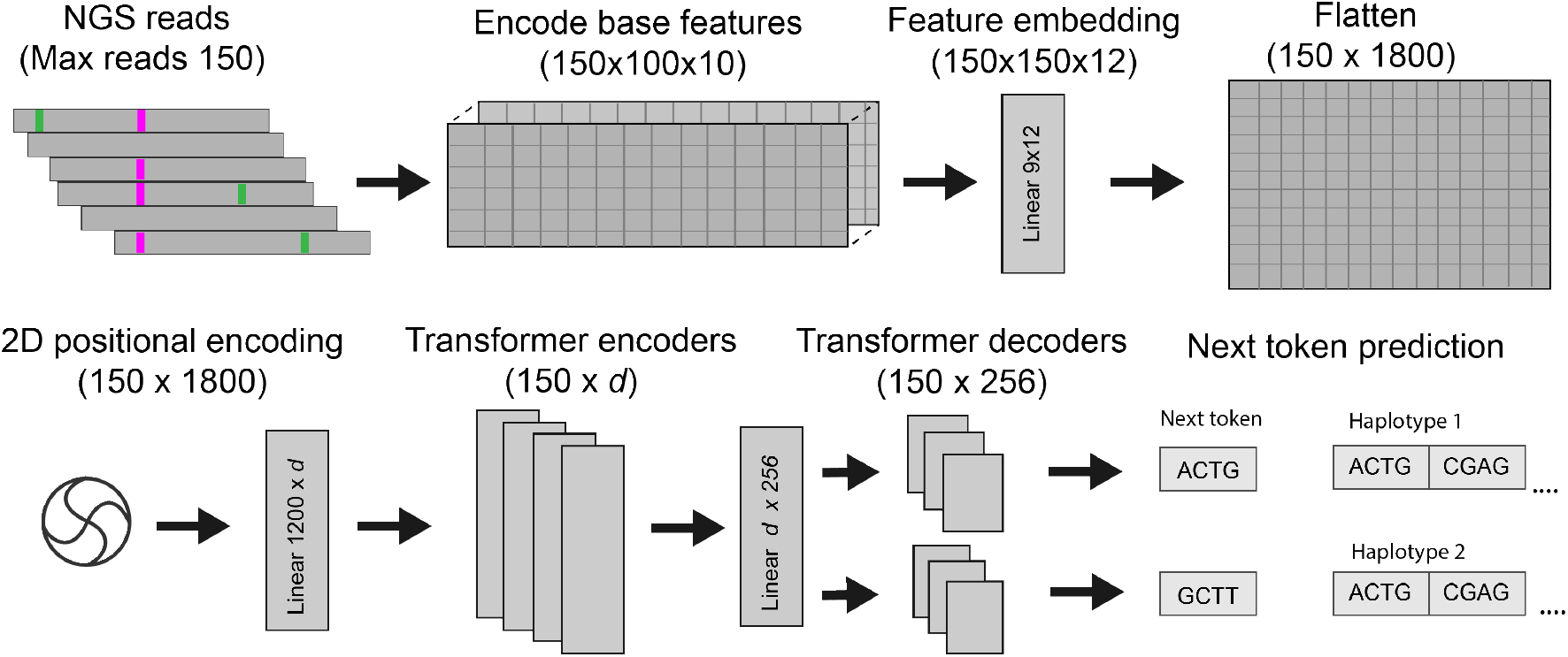
Model architecture for Jenever

### 2.2 Training Data and Procedure

Training data was obtained from 37 Whole Genome Sequences (WGS) from seven Genomein-a-Bottle cell lines. Ten of the samples were prepared with the Illumina TruSeq PCR-free kit, while the others were prepared with the Illumina Nextera DNA Flex kit, which involves several PCR cycles. All samples were sequenced on an Illumina NovaSeq 6000 instrument in 2×150 mode, to an approximate read depth of 50 (range 24.7 - 72.3). After conversion to fastq, the sequenced reads were aligned to human reference genome GRCh37.p13 with the GEM-mapper (v3, Marco-Sola et al. 2012) and were sorted and converted to CRAM format with samtools version 1.9 (Li et al. 2009). No additional refinements, such as duplicate read marking, base-quality score recalibration, or indel realignment were performed.

To select regions to include for training data we developed a scheme to sample regions in a biased manner, prioritizing regions containing variants and, especially, regions with multiple or complex variants. Regions of the reference genome overlapping the high-confidence regions from Genome-in-a-Bottle were subdivided into 150bp windows, and these regions labelled according to the presence of variants. Separate labels were generated for regions containing a single SNV, deletion, or insertion, as well as regions containing multiple insertion-deletions, or those containing variants intersecting low-complexity or porr mappability regions. We additionally included ‘true negative’ regions where no known variant was present. In regions containing more than 150 overlapping reads, the reads were repeatedly downsampled to generate multiple training regions. For all analyses described here, we obtain training data only from the human autosomes 1-20, and hold out chromosomes 21 and 22 for model evaluation. Approximately 1.48M regions were obtained from each of the 37 samples, generating a full training data set of 54.8M regions encompassing 8.2B input tokens.

Target haplotype sequences were produced by obtaining truth variants from the Genomein-a-Bottle VCF files (version 4.2.1, Wagner et al. 2022) for each sample and inserting the variants into the reference sequence. Two sequences were generated for each region, one representing each haplotype. In regions where phasing of the variants was ambiguous, the reads in the sequenced sample were examined to determine phase status. Briefly, all possible genotypes (pairs of haplotypes) were generated and reads were aligned via Smith-Waterman to the possible haplotypes, and the highest scoring genotypes selected as the most likely phasing. This phasing procedure was only attempted for variants less than 100bp apart, otherwise the region was discarded.

Models were trained using the AdamW optimizer with *β* weight decay parameters set to 0.9 and 0.99, as implemented in PyTorch 2.1 (Paszke et al. 2019). We used a learning rate schedule with a linear warm-up from a value near 0 to a maximum of 5 *×* 10^*−*5^, followed by cosine decay to a minimum of 10^*−*5^, and a batch size of 512 regions. For our best-performing model, we performed additional fine-tuning rounds on data with increased representation of poor mappability and segmentally duplicated regions as defined in the Genome-in-a-Bottle stratification files (version 4.2, Wagner et al. 2022). These fine-tuning runs were performed with a learning rate schedule as above, but with a maximum rate of 10^*−*5^ and minimum of 10^*−*6^. Training was performed on 1-2 Nvidia A6000 (Ampere generation) GPUs.

Haplotypes have no intrinsic order, but the model produces two haplotypes that must be matched to the corresponding target haplotypes correctly. If the model produces haplotypes (*A*_*P*_, *B*_*P*_) and the target haplotypes are (*B*_*T*_, *A*_*T*_), we want to compute the loss and corresponding gradients for *L*(*A*_*P*_, *A*_*T*_) + *L*(*B*_*P*_, *B*_*T*_), where *L* is the loss function. For instance, the model shouldn’t be penalized for producing haplotypes with a SNV on haplotype *A* when the targets have the same SNV on haplotype *B*. To ameliorate this, we compute the loss for each haplotype configuration and backpropogate gradients only for the configuration that produces the lowest loss.

### 2.3 Variant detection

Given an alignment file in BAM or CRAM format, we first identify regions where a potential variant might exist. Any genomic position in which at least three reads contain a base that differs from the reference or an indel are flagged as potentially containing a variant. All regions are padded by four basepairs in both directions, and regions closer than 100bp are merged into a single region.

For a single region containing suspected variants, we perform multiple overlapping forward passes of the model with step size *k*, where *k* = 25 for the results reported here, beginning 100 bases upstream of the potential variant. We use the standard greedy decoding procedure to generate haplotypes, selecting the single 4-mer with the highest predicted probability on each iteration until the predicted sequence spans the entire candidate region or until 37 4-mers (148 bases) have been generated. Each haplotype is aligned via Smith-Waterman to the reference sequence, and any mismatching positions are converted to variant calls. After variants are collected across multiple windows results are merged using an ad-hoc method that attempts to minimize the number of conflicting calls between windows using a simple majority-rules algorithm. For instance, if a variant was found to be heterozygous in 2 windows and homozygous in one window, the output variant is predicted to be heterozygous. For each variant we record the number of windows in which each variant was detected, the number of variants in cis and trans, the probability associated with the *k*-mer containing the variant bases, and the position of the variant within each window.

### 2.4 Variant quality calculation

The above variant detection procedure has two shortcomings. First, if every potential variant is emitted, then precision is poor. Second, the model does not innately produce well calibrated variant quality scores. To address these shortcomings we introduce a post-hoc random forest classifier that we train to discriminate true and false-positive variant calls. The resulting score is used as a final variant quality score suitable for tuning the tradeoff between sensitivity and specificity. The random forest classifier was trained by calling variants on selected regions from autosomes 1-2 totalling approximately 90Mb. True and false positive variant calls were identified with the vcfeval tool (Cleary et al. 2015). We used the scikit-learn implementation of the random forest with 100 trees and maximum tree depth of 25. Features are listed in Supplementary Table 3.

## 3 Results

### 3.1 Model size and training data experiments

Overall model accuracy, sensitivity, and precision increased with model size and over the course of training (Figure 2), with most of the gains accuring during the first 3 *×* 10^7^ training regions processed. Precision increased more slowly than sensitivity during training, with raw SNV precision reaching only 80% for the 30M and 50M models after one epoch. Training for additional epochs improved performance somewhat, with SNV precision reaching approximately 98% for the 100M model after 3 epochs. Note that these training evaluation statistics do not reflect the full variant detection procedure outlined in section 2.3, and are derived from a single window without variant quality calculation or filtering. Sensitivity was higher overall and showed less variability across model sizes, although the 100M model consistenly demonstrated a slightly higher SNV accuracy than the smaller models.

**Figure 2:**
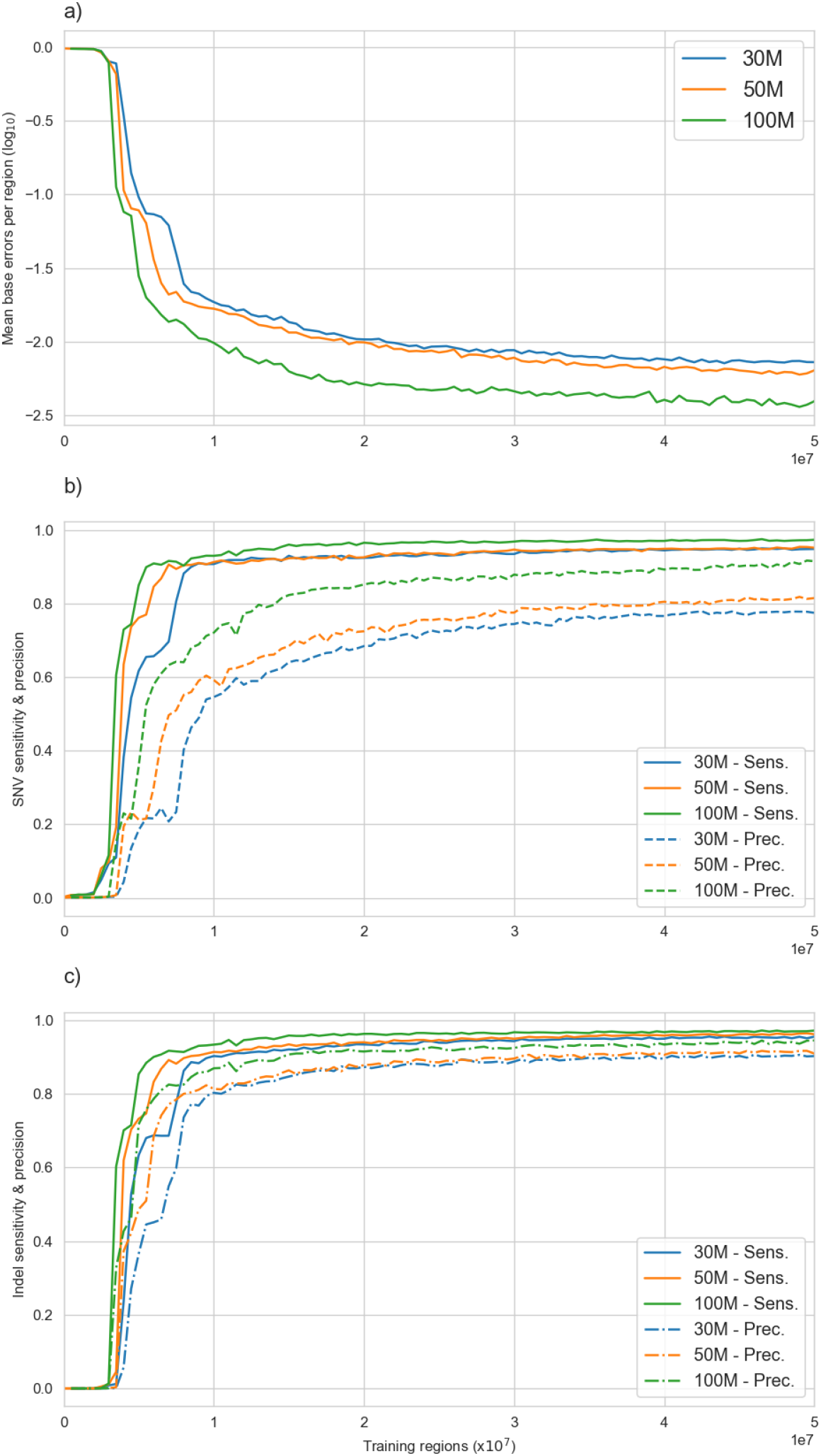
Mean base prediction error rate (a), sensitivity and precision curves for SNVs (b) and indels (b) for the chromosome 21 and 22 validation regions during the first epoch of training for the 30M, 50M, and 100M models. Base error rate is the mean fraction of incorrectly predicted bases in a region. Data8are collected for individual validation regions prior to the multiple window calling procedure and variant quality calculations.

### 3.2 Variant detection accuracy

To characterize variant detection accuracy, we called variants on validation regions from chromosomes 21 and 22 for all 37 samples following the procedure described in section 2.3, and compared results to DeepVariant (v1.3.0, Poplin et al. 2018), GATK HaplotypeCaller (v. 4.1.4.1), FreeBayes 1.3.6 (Garrison & Marth 2012), and Clair3 (Luo et al. 2020). The regions include most of chromosomes 21 and 22, although we omit the p-arm of chromosome 21 which is almost entirely masked on GRCh37, in total comprising about 67.8MB. For this comparison we used our largest model (100M), after training for approximately 3 epochs. We selected filtering criteria for each caller by reviewing ROC curves produced by vcfeval (Cleary et al. 2015) and selecting values close to the *F*_1_ maximizing value across samples. Jenever calls were filtered at quality 10 (phred-scaled), HaplotypeCaller at 50, Clair3 at 0, and DeepVariant at 3. Sensitivity, precision, and related calculations were computed with the *hap*.*py* tool (Krushe et al. 2019) using Genome-in-a-Bottle (v4.2.1) as the benchmark variant set.

Jenever achieved the highest sensitivity and precision for insertion / deletion (indel) calls across all callers we tested, with a mean sensitivity of 98.09% and precision of 98.81% (Figure 3). DeepVariant was the next closest in performance, with a sensitivity of 96.49% and a precision of 98.26%. For single nucleotide variants (SNVs), our model had the highest sensitivity (99.27%) of all callers, but was outperformed by both DeepVariant and Clair3 in precision, with DeepVariant achieving the highest overall precision of 99.89% and Jenever yielding 99.56%. Overall, our model had the highest *F*_1_ score for both indels (98.45%) and SNVs (99.42%) (Figure 4), surpassing the nearest competitor, DeepVariant, by more than a full percentage point (1.08%) for indels and 0.3% for SNVs. Jenever showed especially high accuracy for small (1-5bp) to medium size (6-15bp) insertions and deletions, and both insertion and deletion variants had similar sensitivity and precision. For large deletions (*>*15bp) Jenever had somewhat lower precision than other callers, but similar sensitivity, while Jenever demonstrated high sensitivity and average precision for insertions *>* 15bp.

**Figure 3:**
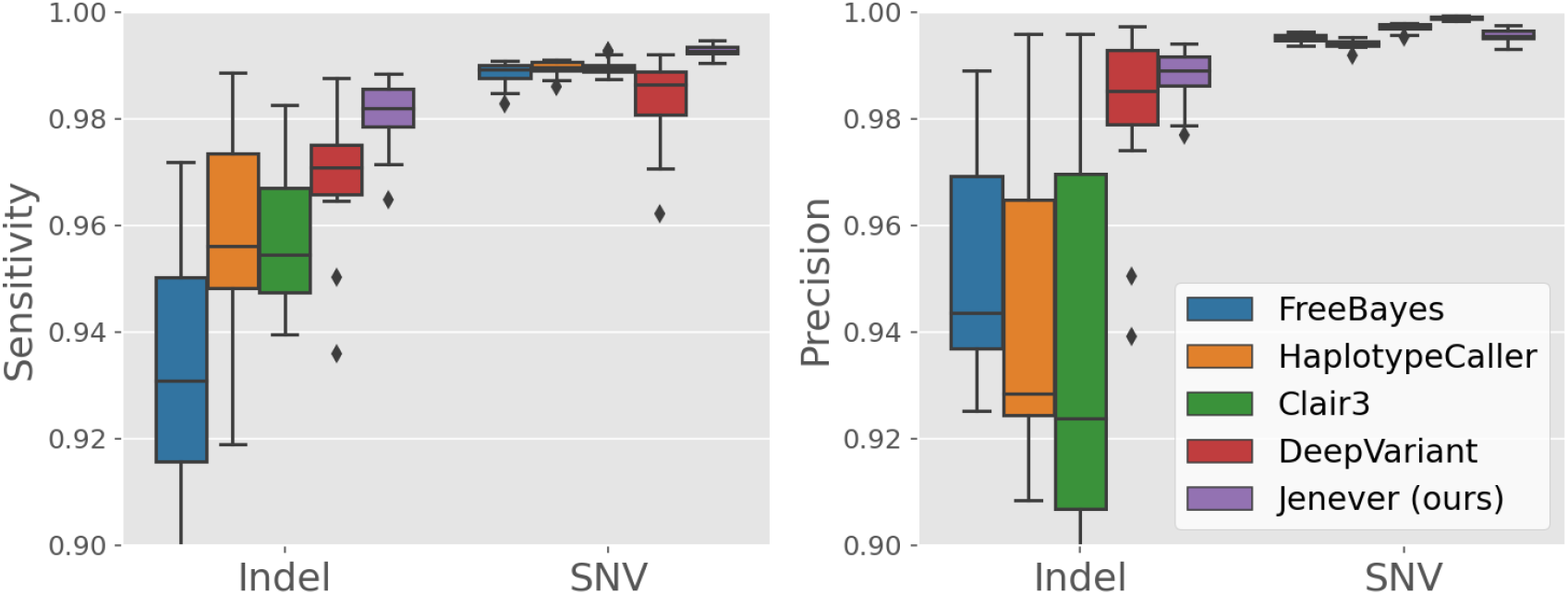
Mean per-sample sensitivity (a) and precision (b) for indels and SNVs in validation regions as computed by *hap*.*py* across variant callers. Error bars denote 95% confidence intervals.

**Figure 4:**
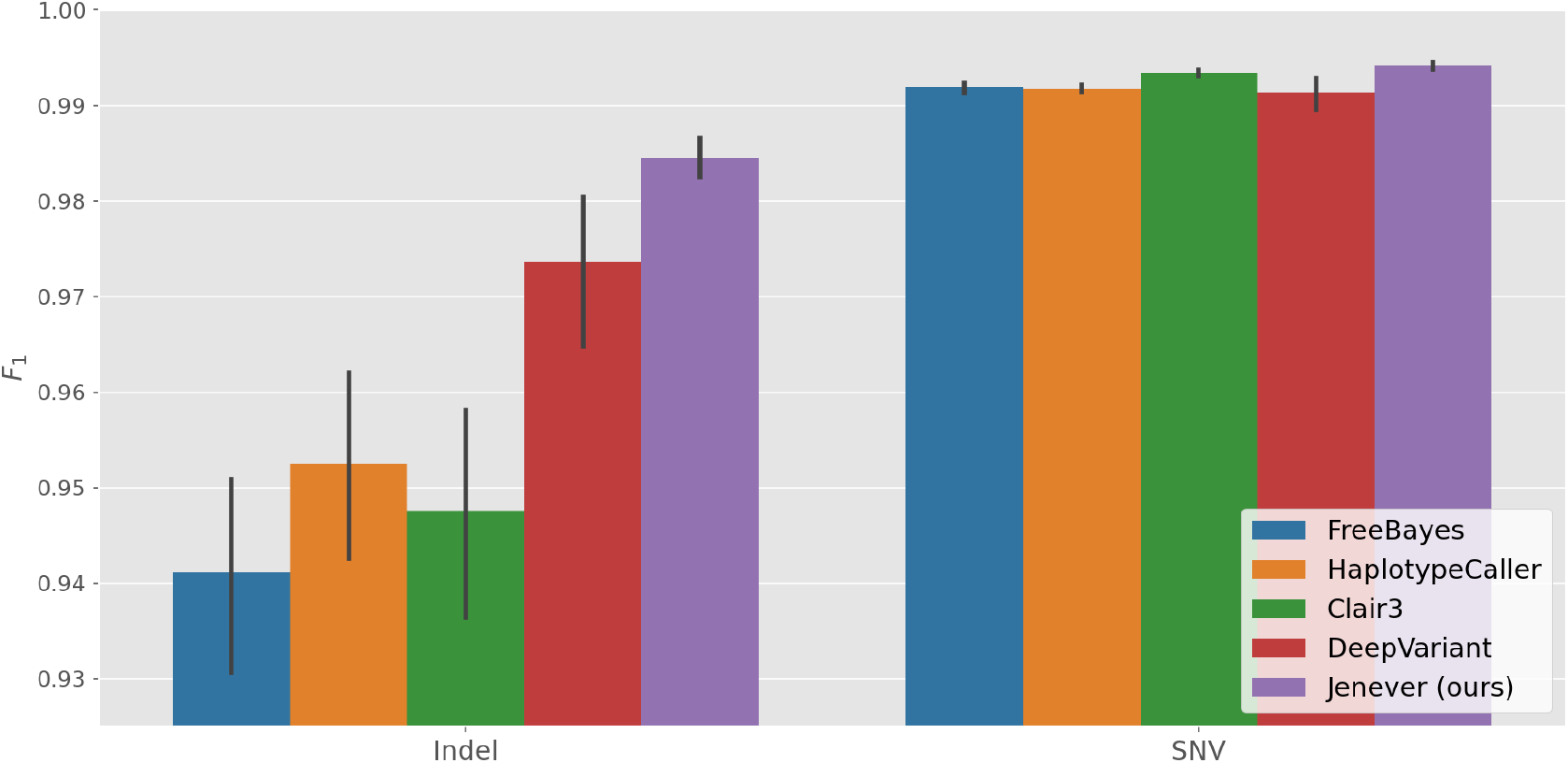
Mean per-sample *F*_1_ scores for variants in validation regions, as computed by *hap*.*py* for all variant callers examined. Error bars denote 95% confidence intervals.

**Figure 5:**
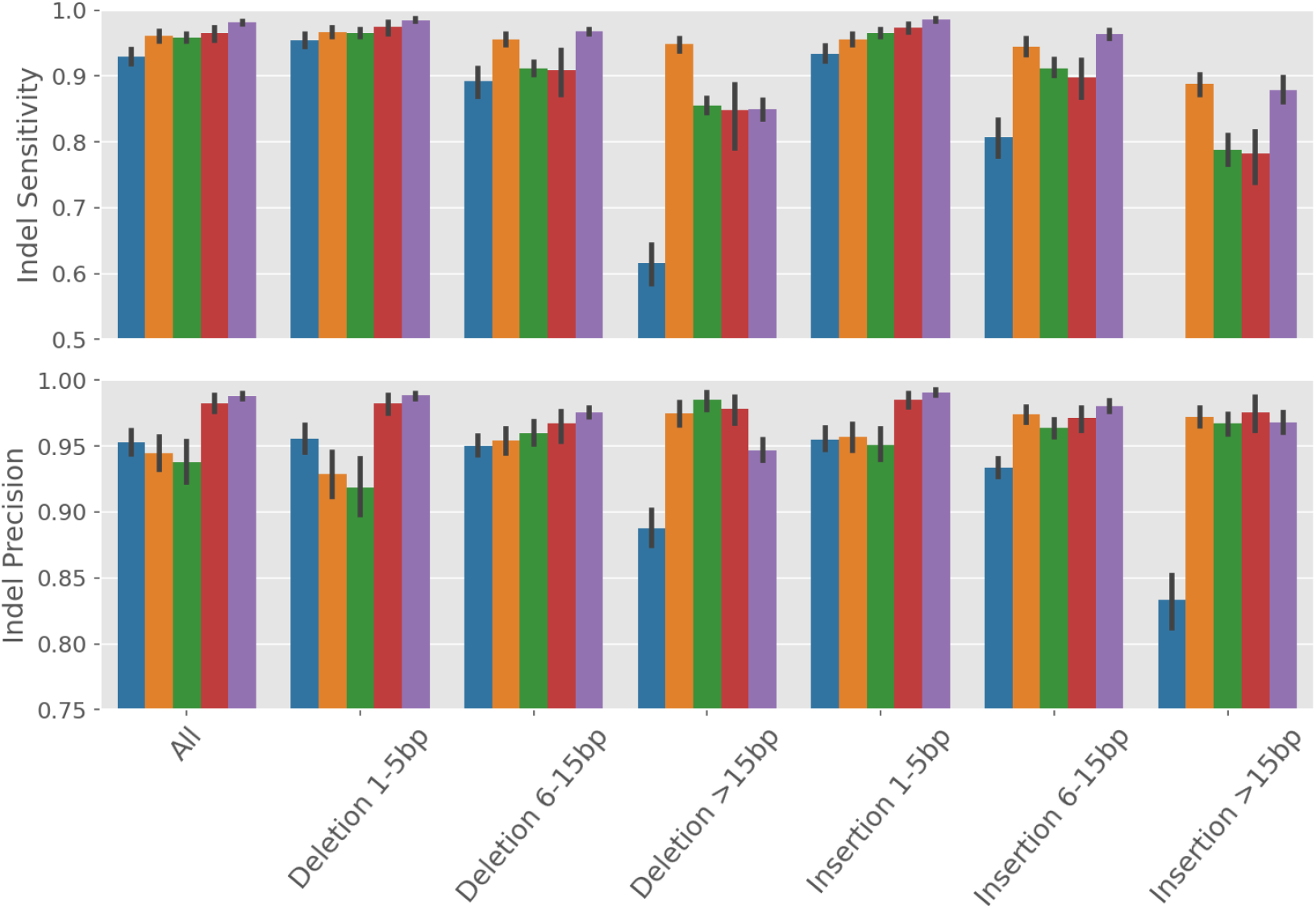
Sensitivity and precision for insertion and deletion variants binned by size. Error bars denote 95% confidence intervals.

### 3.3 Genotype & phasing accuracy

A key feature of our model is the use of two parallel transformer decoder blocks to generate haplotypes. The decoder blocks do not commununicate, leaving the encoder block as the only mechanism to ’decide’ which haplotype each decoder must generate. To assess the accuracy of this system we examine both the genotype predictions of individual calls and the phase predictions for pairs of nearby calls by comparing them to Genome-in-a-Bottle reference data.

We examined genotype accuracy by tallying the number of variants in validation regions where the callers produced the correct allele and position, but the incorrect genotype, as computed by *hap*.*py*. Jenever produced the fewest incorrect genotype calls for indel variants (mean 83.4 per sample), but the most incorrect genotypes for SNVs (mean 93.8).

By explicitly modeling haplotypes, our method provides a form of read-backed phasing. We investigate the accuracy of our phase predictions by comparing them to data from Genome-in-a-Bottle reference sample HG002, the only reference sample for which phasing data is provided. Because the reference used pedigree information to phase the variants we cannot match most of the predictions, and instead we assess how often the phase predictions from our model match those from the reference sample. Specifically, we find groups of phased variants in our predictions, and for each adjacent pair of variants in the same phase group we compare to the reference sample. Occurences in which our phase predictions matched or mismatched those from the reference were tallied, and cases where no phase information was provided from the reference, or variant genotype was found to conflict with the reference, were discarded. For this analysis we discarded variants with a phred-scaled quality score of less than 10, in a manner identical to the previous section.

Phase accuracy was high overall and decreased with distance between variants (Table 2).

**Table 1:** Model parameter combinations used for the three model sizes investigated.

| Model name | Encoder layers | Encoder heads | Embedding dim. | Decoder layers | Decoder heads |
| --- | --- | --- | --- | --- | --- |
| 30M | 6 | 8 | 800 | 4 | 4 |
| 50M | 8 | 8 | 960 | 8 | 4 |
| 100M | 10 | 8 | 1280 | 10 | 10 |

**Table 2:**
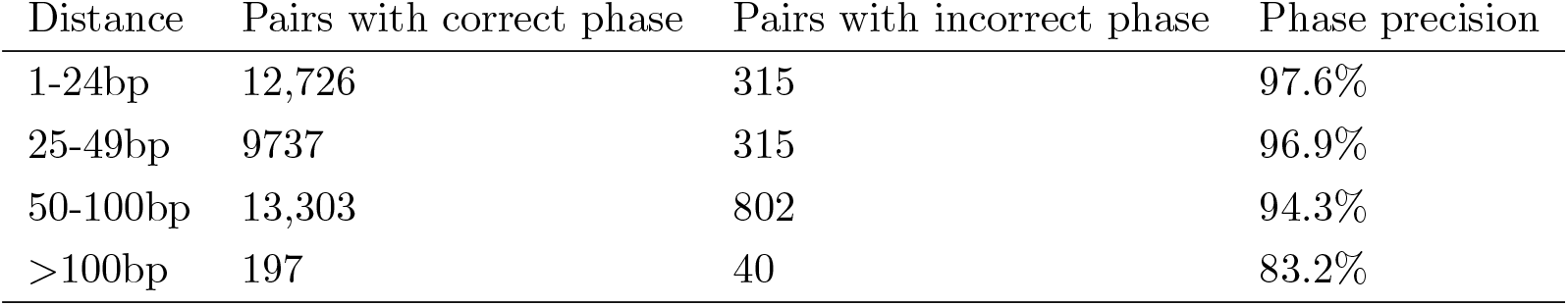
Accuracy of phase predictions compared to reference sample HG002 for pairs of variants in chromosome 21-22 validation regions.

**Table 3:** Features used in the variant quality prediction model.

| Feature | Description |
| --- | --- |
| Mean probability | Mean probability associated with first token containing variant across windows |
| Ref length | Length of reference allele |
| Alt length | Length of longest alternate allele |
| Min probability | Minimum probability of first token containing variant across windows |
| Max probability | Maximum probability of first token containing variant across windows |
| Window var count | Total number of variants in all windows |
| Cis count | Total number of variants on same haplotype as this variant |
| Trans count | Total number of variants on opposite haplotype as this variant |
| Step count | Total number of windows in which this variant was called |
| Window count | Total number of windows overlapping this variant position |
| Min window offset | Minimum distance from variant position to start of calling window |
| Max window offset | Maximum distance from variant position to start of calling window |
| Depth | Total number of reads overlapping this variant position |
| Genotype | Heterozygous or homozygous |
| Variant allele frequency | Fraction of reads containing this variant allele |
| Read support | Total number of reads containing this variant allele |

Over 95% of variant pairs has phase predictions that matched the reference sample when variants were within 50bp, but accuracy drops to just over 80% for variants that are greater than 100bp apart. Anecdotally, most incorrect predictions appear in pairs involving indels in low complexity sequence.

## 4 Discussion

We describe a new approach to detecting sequence variants in short-read NGS data that utilizes a deep generative model of haplotypes. Our model takes short NGS reads aligned to the reference genome as input, and learns to predict the two germline haplotypes present in the sample. The model architecture is very similar to that of modern large language models (LLMs), but we replace the input ”prompt” with aligned NGS reads, and instead of generating words we generate nucleotide *k*-mers representing haplotypes. The advantage of this approach is that the model fully encompasses both the candidate allele generation and evaluation components, and thus and can be optimized directly to produce accurate haplotypes. In contrast to other common variant detection tools, our approach does not require any handcrafted statistical methods, bespoke algorithms, or finely-tuned thresholds.

Our model is significantly more accurate than existing small variant detectors, especially for insertion and deletion variants. At quality thresholds chosen to maximize the *F*_1_ statistic, our model delivers a mean sensitivity more than 1.5% higher than the next closest model (98.0%, compared to 96.5% for DeepVariant) at improved precision (98.8%, compared to 98.2% for DeepVariant). Additionally, our model makes half as many genotyping errors for indels (variants where the correct allele was detected but with the wrong genotype) as the next closest competitor (Figure 6). For single nucleotide variants (SNVs) results are more mixed. Both DeepVariant and Clair3 offer higher precision at *F*_1_ maximizing quality thresholds, but at the cost of significantly reduced sensitivity. Our model produces more false positive calls and more genotyping errors, but detects many additional true positive calls that are missed by other callers, leading to a higher *F*_1_ accuracy overall. Jenever’s mean precision for SNVs was 99.5% while DeepVariant delivered an impressive 99.9%, however DeepVariant’s sensitivity was only 98.4%, while our model found 99.3% of real variants.

**Figure 6:**
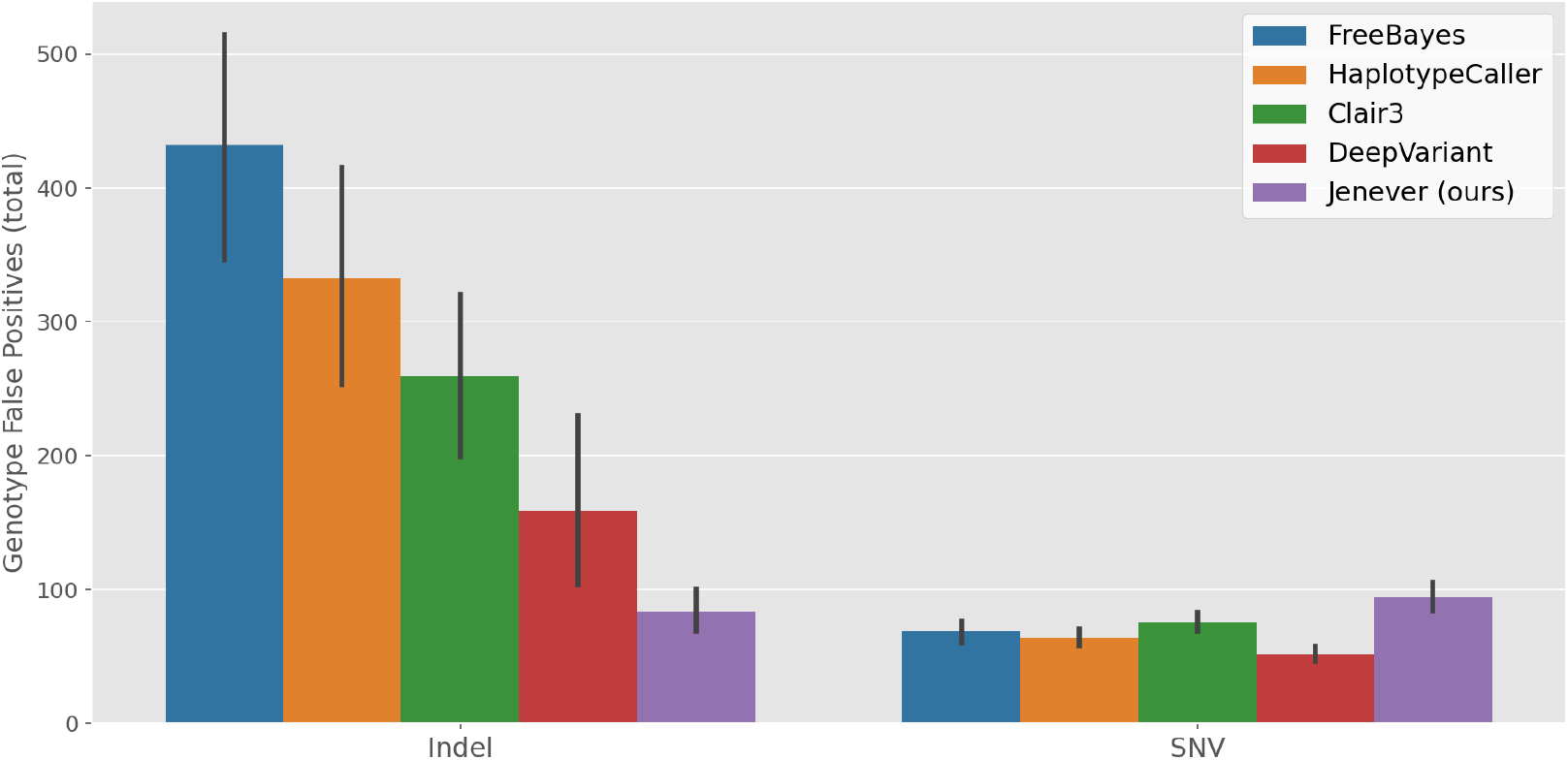
Mean number of incorrect genotype calls for indels and SNVs in validation regions for all callers examined.

We hypothesize that the performance difference between indels and SNVs in our model may be due to the attention mechanism used in the transformer architecture. Both the self- and cross-attention mechanisms in the transformer architecture were originally designed for language modeling (Vaswani et al. 2017) and rely on computing the interaction between tokens, where tokens represent information across genomics positions. Intuitively, this seems well-suited to inferring indel variants, where the model must utilize information from multiple nearby tokens to produce the correct output. In contrast, SNV detection typically requires utilizing information from a single genomic position. Manually reviewing some of the incorrect SNV calls suggests that Jenever struggles in differentiating true and false calls at lower allele frequencies, a task where multi-token attention is less useful. In earlier work, we demonstrated that a similar model used the number of nearby reference mismatches to inform SNV detection for a given site (O’Fallon et al. 2022), suggesting that the model can in some cases employ information from the local context to improve SNV detection, but this mechanism appears less accurate than convolutional approaches used in other modern models.

While our implementation is very accurate, it is much slower than other tools. Generation of each output token (4-mer) requires a separate pass through both haplotype decoders, each of which is a deep neural net with tens of millions of parameters, and each candidate region requires multiple passes of the model for high prediction accuracy. Despite this, numerous avenues exist for improving runtime performance, including model quantization, pruning, distillation, multi-query attention (Shazeer 2019), and many others. In general, improving transformer inference speeds is a topic of intense interest in both the academic and business communities, with recent significant advances in both software (e.g. Dao et al. 2022) and hardware (e.g. Qi et al. 2021). We plan to explore methods for improving runtime performance in future work.

By framing NGS variant detection as a transformer-based deep generative model, we are able to leverage the many recent advances in transformer training and inference optimizations. For instance, improvements in positional embeddings (e.g. Shaw et al. 2019, Su et al. 2024), attention mechansisms (e.g. Choromanski et al. 2020, Roy et al. 2021), ensembling methods (e.g. Riquelme et al. 2021) or training methodologies (e.g. Izmailov et al. 2018), although originally designed for language or vision transformers, are likely to provide benefits for our model as well. By combining these and other advances numerous opportunities exist for delivering the next generation of variant callers.

## 5 Data and Code Availability

All source code and training scripts are available via git at https://github.com/brendanofallon/jovian. Model weights and working example are provided via Dockerhub at [to be determined upon publication].

## 7 Supplementary Tables

## References

Baid, Gunjan, et al. “DeepConsensus improves the accuracy of sequences with a gap-aware sequence transformer.” Nature Biotechnology 41.2 (2023): 232–238.

Behera, Sairam, et al. “Comprehensive and accurate genome analysis at scale using DRAGEN accelerated algorithms.” bioRxiv (2024): 2024–01.

Choromanski, Krzysztof, et al. “Rethinking attention with performers.” arXiv preprint arXiv:2009.14794 (2020).

Cleary, John G., et al. “Comparing variant call files for performance benchmarking of next-generation sequencing variant calling pipelines.” BioRxiv (2015): 023754.

Cooke, Daniel P., David C. Wedge, and Gerton Lunter. “A unified haplotype-based method for accurate and comprehensive variant calling.” Nature biotechnology 39.7 (2021): 885–892.

Dao, Tri, et al. “Flashattention: Fast and memory-efficient exact attention with io-awareness.” Advances in Neural Information Processing Systems 35 (2022): 16344–16359.

DePristo, Mark A., et al. “A framework for variation discovery and genotyping using next-generation DNA sequencing data.” Nature genetics 43.5 (2011): 491–498.

Dosovitskiy, Alexey, et al. “An image is worth 16×16 words: Transformers for image recognition at scale.” arXiv preprint arXiv:2010.11929 (2020).

Garrison, Erik, and Gabor Marth. “Haplotype-based variant detection from short-read sequencing.” arXiv preprint arXiv:1207.3907 (2012).

Goldfeder, Rachel L., et al. “Medical implications of technical accuracy in genome sequencing.” Genome medicine 8.1 (2016): 1–12.

Gupta, Gaurav, and Shubhi Saini. “DAVI: Deep learning-based tool for alignment and single nucleotide variant identification.” Machine Learning: Science and Technology 1.2 (2020): 025013

Kim, Sangtae, et al. “Strelka2: fast and accurate calling of germline and somatic variants.” Nature methods 15.8 (2018): 591–594.

Krusche, Peter, et al. “Best practices for benchmarking germline small-variant calls in human genomes.” Nature biotechnology 37.5 (2019): 555–560.

Li, Heng. “Toward better understanding of artifacts in variant calling from high-coverage samples.” Bioinformatics 30.20 (2014): 2843–2851.

Li, Heng, et al. “The sequence alignment/map format and SAMtools.” Bioinformatics 25.16 (2009): 2078–2079.

Li, Heng, and Richard Durbin. “Fast and accurate short read alignment with Burrows–Wheeler transform.” Bioinformatics 25.14 (2009): 1754–1760.

Liu, Ze, et al. “Swin transformer: Hierarchical vision transformer using shifted windows.” Proceedings of the IEEE/CVF International Conference on Computer Vision. 2021.

Luo, Ruibang, et al. “A multi-task convolutional deep neural network for variant calling in single molecule sequencing.” Nature communications 10.1 (2019): 1–11.

Luo, Ruibang, et al. “Exploring the limit of using a deep neural network on pileup data for germline variant calling.” Nature Machine Intelligence 2.4 (2020): 220–227.

Marco-Sola S., Sammeth M., Guigó R., Ribeca P. “The GEM mapper: fast, accurate and versatile alignment by filtration”. Nat Methods. (2012);9(12):1185–1188. doi:10.1038/nmeth.2221

Nielsen, Rasmus, et al. “Genotype and SNP calling from next-generation sequencing data.” Nature Reviews Genetics 12.6 (2011): 443–451.

O’Fallon, Brendan, et al. “Jovian enables direct inference of germline haplotypes from short reads via sequence-to-sequence modeling.” bioRxiv (2022): 2022–09.

Paszke, A., et al. “PyTorch: An Imperative Style, High-Performance Deep Learning Library.” In Advances in Neural Information Processing Systems 32 (2019):8024–8035. Curran Associates, Inc. Retrieved from http://papers.neurips.cc/paper/9015-pytorch-an-imperative-style-high-performance-deep-learning-library.pdf

Petti, Samantha, et al. “End-to-end learning of multiple sequence alignments with differentiable Smith-Waterman.” BioRxiv (2021).

Poplin, Ryan, et al. “A universal SNP and small-indel variant caller using deep neural networks.” Nature biotechnology 36.10 (2018): 983–987.

Poplin, Ryan, et al. “Scaling accurate genetic variant discovery to tens of thousands of samples.” BioRxiv (2018): 201178.

Qi, Panjie, et al. “Accelerating framework of transformer by hardware design and model compression co-optimization.” 2021 IEEE/ACM International Conference On Computer Aided Design (ICCAD). IEEE, 2021.

Ramachandran, Anand, et al. “HELLO: A hybrid variant calling approach.” bioRxiv (2020).

Riquelme, Carlos, et al. “Scaling vision with sparse mixture of experts.” Advances in Neural Information Processing Systems 34 (2021): 8583–8595.

Roy, Aurko, et al. “Efficient content-based sparse attention with routing transformers.” Transactions of the Association for Computational Linguistics 9 (2021): 53–68.

Shaw, Peter, Jakob Uszkoreit, and Ashish Vaswani. “Self-attention with relative position representations.” arXiv preprint arXiv:1803.02155 (2018).

Shazeer, Noam. “Fast transformer decoding: One write-head is all you need.” arXiv preprint arXiv:1911.02150 (2019).

Su, Jianlin, et al. “Roformer: Enhanced transformer with rotary position embedding.” Neurocomputing 568 (2024): 127063.

Izmailov, Pavel, et al. “Averaging weights leads to wider optima and better generalization.” arXiv preprint arXiv:1803.05407 (2018).

Vaswani, Ashish, et al. “Attention is all you need.” Advances in neural information processing systems 30 (2017).

Wagner, Justin, et al. “Benchmarking challenging small variants with linked and long reads.” Cell Genomics 2.5 (2022).

